# A potential role for GrgA in regulation of σ^28^-dependent transcription in the obligate intracellular bacterial pathogen *Chlamydia trachomatis*

**DOI:** 10.1101/322701

**Authors:** Malhar Desai, Wurihan Wurihan, Rong Di, Joseph D. Fondell, Bryce E. Nickels, Xiaofeng Bao, Huizhou Fan

**Affiliations:** Department of Pharmacology, Robert Wood Johnson Medical School, Rutgers University, Piscataway,10 NJ 08854; Graduate Program in Physiology and Integrative Biology, School of Graduate Studies, Rutgers University, NJ 08854; Department of Plant Biology, School of Environmental and Biological Sciences, Rutgers University, New Brunswick, NJ 08901; Waxman Institute of Microbiology, Rutgers University, Piscataway, NJ 08854; Department of Pharmacology, School of Pharmacy, Nantong University, Nantong, China 226001

**Author notes:** Address correspondence to Huizhou Fan or Xiaofeng Bao.

## Abstract

The sexually transmitted obligate intracellular bacterial pathogen *Chlamydia trachomatis* has a unique developmental cycle consisting of two contrasting cellular forms. Whereas the primary *Chlamydia* sigma factor, σ^66^, is involved in the expression of the majority of chlamydial genes throughout the developmental cycle, expression of several late genes requires the alternative sigma factor σ^28^. In prior work we identified GrgA as a *Chlamydia-*specific transcription factor that activates σ^66^-dependent transcription by binding DNA and interacting with a non-conserved region (NCR) of σ^66^. Here, we extend these findings by showing GrgA can also activate σ^28^-dependent transcription through direct interaction with σ^28^. We measure the binding affinity of GrgA for both σ^66^and σ^28^, and we identify regions of GrgA important for σ^28^-dependent transcription. Similar to results obtained with σ^66^, we find that GrgA’s interaction with σ^28^ involves a NCR located upstream of conserved region 2 of σ^28^. Our findings suggest GrgA is an important regulator of both σ^66^- and σ^28^-dependent transcription in *C. trachomatis* and further highlight NCRs of bacterial RNA polymerase as targets for regulatory factors unique to particular organisms.

## IMPORTANCE

*Chlamydia trachomatis* is the number one sexually transmitted bacterial pathogen worldwide. A substantial proportion of *C. trachomatis*-infected women develop infertility, pelvic inflammatory syndrome and other serious complications. *C. trachomatis* is also a leading infectious cause of blindness in under-developed countries. The pathogen has a unique developmental cycle, which is transcriptionally regulated. The discovery of an expanded role for the *Chlamydia*-specific transcription factor GrgA helps understand progression of the chlamydial developmental cycle.

## INTRODUCTION

Each year, about 2.2 million cases of notifiable infections are reported to the Centers for Disease Control and Prevention (CDC). These infections are caused by nearly 100 different pathogens, but the majority (about 1.6 million, i.e., 60%) is due to the sexually transmitted pathogen *Chlamydia trachomatis* (1, 2). Still, CDC estimates that only 1 tenth of *C. trachomatis*-infected cases are reported because the infection is mostly asymptomatic (3). Nonetheless, without proper antibiotic treatment, the infection often leads to serious complications. In fact, *C. trachomatis* is the most common infectious cause of infertility and pelvic inflammatory syndrome in women. Infection in pregnant women may result in abortion or premature birth. Pathological changes in the fallopian tubes caused by *C. trachomatis* infection may lead to ectopic pregnancy, which causes severe bleeding and likely death if the ectopically embedded embryo is not detected and terminated early enough. Infants may develop *C. trachomatis* pneumonia following acquisition of the pathogen while passing the birth canal of an infected mother. Some *C. trachomatis* serotypes cause ocular infection, and are still the most common infectious microbes associated with blindness in underdeveloped countries (4, 5).

Like other chlamydiae, *C. trachomatis* is an obligate intracellular Gram-negative bacterium that exists in two cellular forms with contrasting properties (6). The small elementary body (EB) is infectious and capable of extracellular survival, but incapable of proliferation. Following binding to a cellular receptor(s), the EB enters a host cell membrane-derived vacuole through endocytosis (7). Within the vacuole termed inclusion, the EB differentiates into a larger cellular form termed reticulate body (RB) within several h. No longer infectious, the RB divides exponentially by binary fission until around 20 h when a significant portion of RBs re-differentiate back into EBs while some RBs continue proliferation (8). Progeny EBs along with residual RBs are released from infected cells following cell lysis. Alternatively, whole inclusions may be released from infected cells (9).

The 1 million bp *C. trachomatis* genome encodes fewer than 1000 genes (10). Microarray analyses demonstrated that the majority of these genes are transcribed starting a few hours post-inoculation throughout the remaining developmental cycle, whereas a small number of genes are transcribed immediately following cell entry and another small set of genes are transcribed only at late stages (11, 12). RNA-seq detected distinct sets of gene transcripts specifically enriched in either EBs or RBs (13), and purified EBs and RBs have been found to transcribe different sets of genes in axenic media (14). These findings suggest that the progression of the chlamydial developmental cycle is transcriptionally regulated.

Transcription is initiated following binding of the RNA polymerase (RNAP) to the gene promoter (15). The bacterial RNAP holoenzyme is comprised of the catalytic core enzyme and a σ factor, which is required for promoter recognition (16). Transcription of the vast majority of *C. trachomatis* genes involves σ^66^, a homolog of σ^70^ that is often referred as the housekeeping σ factor in eubacteria (16). Expression of some (but not all) chlamydial late genes depends on σ^28^. Several genes possess both a σ^66^ promoter and a σ^28^ promoter (17).

GrgA (with the gene codes CT_504 and CTL0766 for *C. trachomatis* serovar D and L2, respectively) is a *Chlamydia-specific* transcription activator (18). It was identified as a protein bound to the σ^66^-dependent promoter of *def*A, which encodes peptide deformylase, an enzyme required for bacterial protein maturation and regulated protein degradation. In addition to *def*A, a midcycle gene, GrgA also stimulates transcription from another midcycle promoter (*omp*A), an early promoter (rRNA P1) and a late promoter (*hct*A), suggesting that GrgA functions as a general activator of σ^66^-dependent genes (18). In this report, we demonstrate that GrgA also stimulates σ^28^-dependent gene transcription *in vitro*. Thus, our findings suggest GrgA plays an expanded role in gene expression during the *C. trachomatis* developmental cycle as a regulator of both σ^66^- and σ^28^-dependent transcription.

## RESULTS

### GrgA physically interacts with σ^28^

To assess whether GrgA potentially regulates expression of σ^28^-dependent genes, we determined whether GrgA can interact with σ^28^. We performed protein pull-down assays using differential epitope-tagging. The StrepTactin beads, which have affinity for the strep tag (19), precipitated NH-σ^28^ (N-terminally poly-His-tagged *C. trachomatis* σ^28^) in a manner that was dependent on the N-terminally strep-tagged GrgA (NS-GrgA) (Fig. 1A).Reciprocally, NH-GrgA was pulled down in an NS-σ^28^-dependent manner (Fig. 1B). The results establish that GrgA can directly interact with σ^28^.

**Fig. 1.**
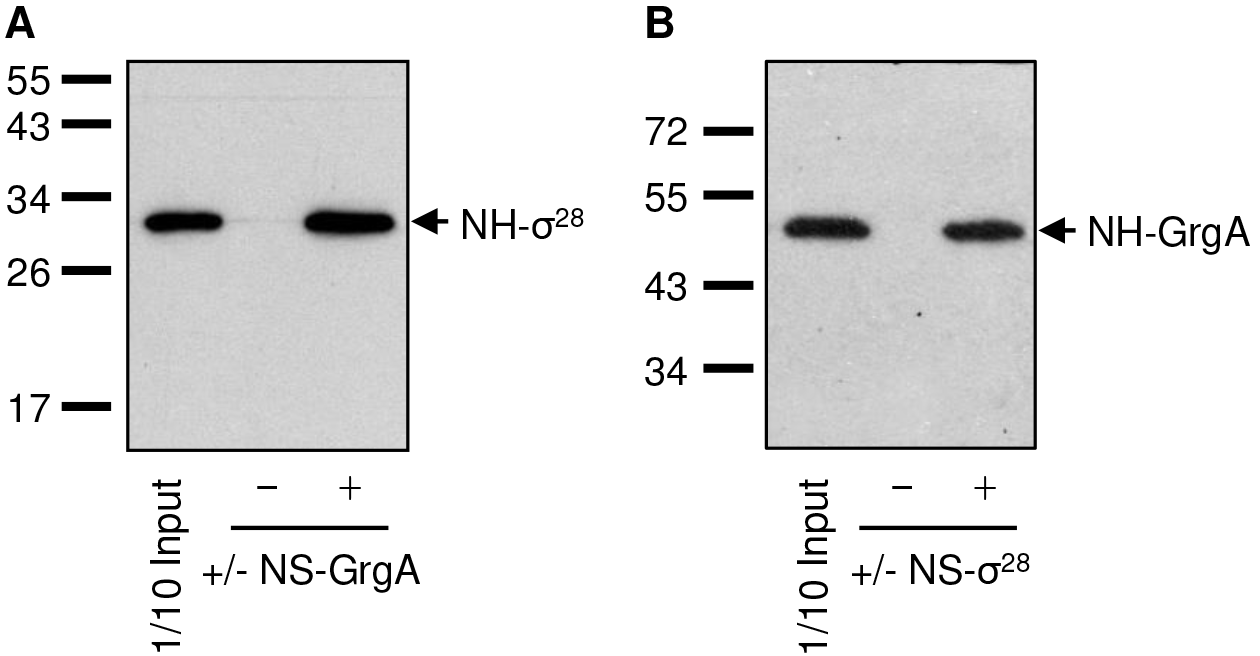
GrgA physically interacts with σ^28^. (A) Pull-down of NH-σ^28^ by StrepTactin bead-immobilized NS-GrgA. Shown isa western blot detecting NH-σ^28^. (B) Pull-down of NH-GrgA by StrepTactin bead-immobilized NS-σ^28^. Shown is a western blot detecting NH-GrgA.

### GrgA has a lower affinity for σ^28^ than for σ^66^

Next, we determined the binding affinities of GrgA for both σ^28^ and σ^66^. We first compared the efficiencies of the σ factors in GrgA-binding by performing competitive pull-down assays. As expected, NS-GrgA efficiently pulled down NH-σ^28^ and CH-σ^66^ in separate reactions (Fig. S1). However, in the presence of equal molar concentrations of NH-σ^28^ and CH-σ^66^, NS-GrgA pulled down only CH-σ^66^ but not NH-σ^28^ (Fig. S1), indicating that GrgA has a lower affinity for σ^28^ than σ^66^.

We next quantitatively characterized GrgA-binding by σ^28^ and σ^66^ with biolayer interferometry using the BLItz system, which detects light wavelength shifts at the biosensor tip with an immobilized ligand following binding of an analyte in a real-time manner (20). Whereas representative BLItz recordings using NH-GrgA as a ligand, and CS-σ^66^ and NS-σ^28^ an analyte are shown in Fig. S2A, B), values of kinetic parameters are provided in Table 1. The CS-σ^66^ analyte yielded a statistically highly significant 25-fold higher *k*_a_ than the NS-σ^28^ analyte, suggesting that CS-σ^66^ binds NH-GrgA much faster than NS-σ^28^. CS-σ^66^ also demonstrated a 3-fold statistically significant increase in *k*_d_, suggestive of moderately higher dissociation from NH-GrgA. Compared to the NH-GrgA-CS-σ^66^ interaction, the NH-GrgA-NS-σ^28^ interaction had a 32-fold higher KD, indicating that GrgA has a lower overall affinity for σ^28^ than σ^66^.

**Table 1.**
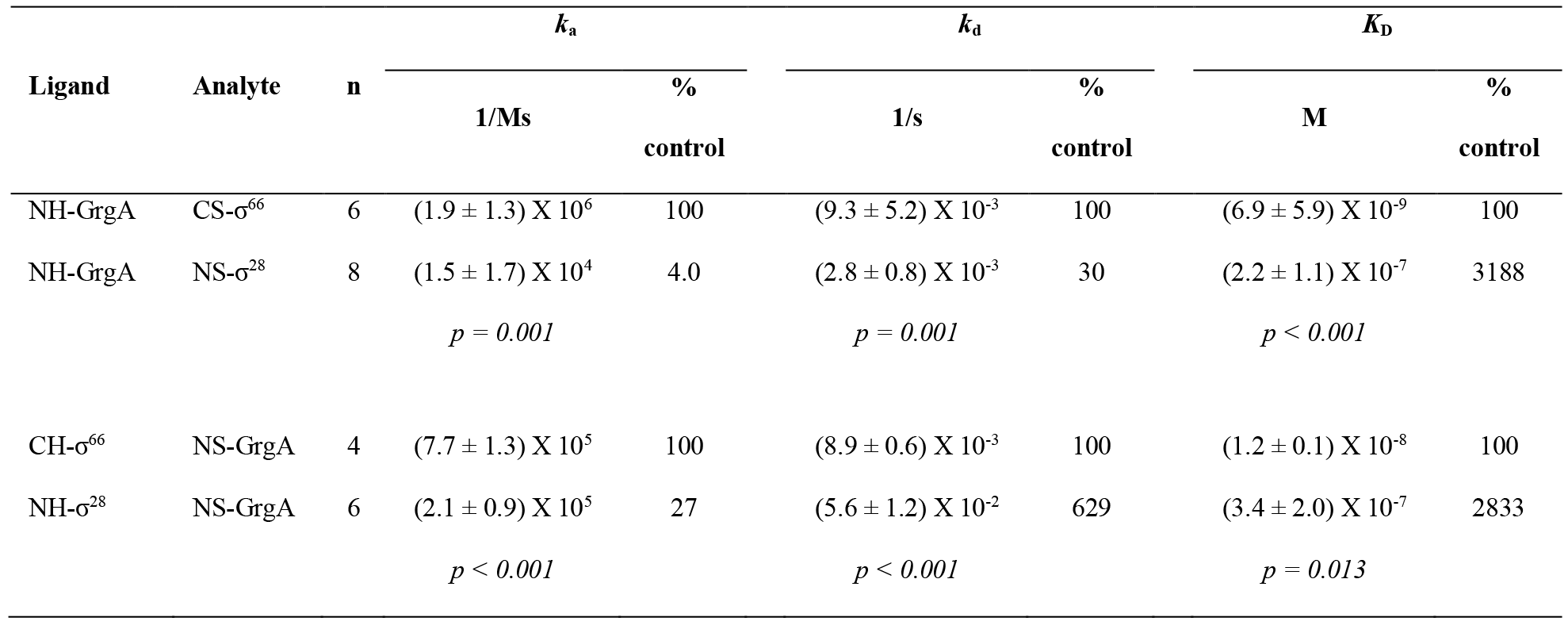
GrgA binds σ^66^ with a higher affinity than σ^28^. BLItz assays were performed with His biosensors using indicated ligand and analyte pairs. Representative graphs of recordings are shown in Fig. S2. Values of kinetic parameters (averages ± standard deviations) were generated by the BLItz Pro software (20). *k*_a_ (association rate constant) is defined as the number of complexes formed per s in a 1 molar solution of A and B. *k*_d_ (disassociation rate constant) is defined as the number of complexes that decay per second. *k*_D_ (disassociation equilibrium constant), defined as the concentration at which 50% of ligand binding sites are occupied by the analytes, is *k*_d_ divided by *k*_a_. n, number of experimental repeats. *p* values were calculated using 2-tailed Student’s *t* tests.

Reciprocal BLI using CH-σ^66^ and NH-σ^28^ as ligands and NS-GrgA as the analyte were performed to validate the difference in GrgA binding by the σ factors presented above (Fig. S2C, D and Table 1). Consistent with the trend in *k*_a_ value changes presented above, the NS-GrgA analyte also demonstrated a statistically significant higher *k*_a_ for CH-σ^66^ than for NH-σ^28^ although the difference is smaller (25-fold vs 3.7 fold). Interestingly, the *k*_d_ values reveal that NS-GrgA also dissociates from CH-σ^66^ 6-fold slower than from NH-σ^28^. Compared to the CH-σ^66^-NS-GrgA interaction, the NH-σ^28^-NS-GrgA interaction had a 28fold higher *K*_D_, which is nearly identical to the 32-fold higher *K*_D_ detected for the NH-GrgA-NS-σ^28^ interaction vs the NH-GrgA-CS-σ^66^ interaction. Thus, competitive pull-down assays and BLI establish that GrgA has a lower affinity for σ^28^ than for σ^66^.

### GrgA stimulates σ^28^-dependent transcription

To determine whether GrgA can stimulate σ^28^-dependent transcription, we performed *in vitro* transcription assays using pMT1212, a transcription reporter plasmid carrying the promoter of a gene encoding a histone-like protein (*hct*B) in *C. trachomatis* (21). Consistent with previous findings (21), transcription from the *hctB* promoter required the addition of NH-σ^28^ to the *C. trachomatis* RNAP (Fig. 2A). Interestingly, GrgA demonstrated a dose-dependent stimulatory effect on the transcription from the promoter (Fig. 2B, C). These data suggest that GrgA can increase the expression of genes with a σ^28^-dependent promoter in addition to genes with a σ^66^-dependent promoter.

**Fig. 2.**
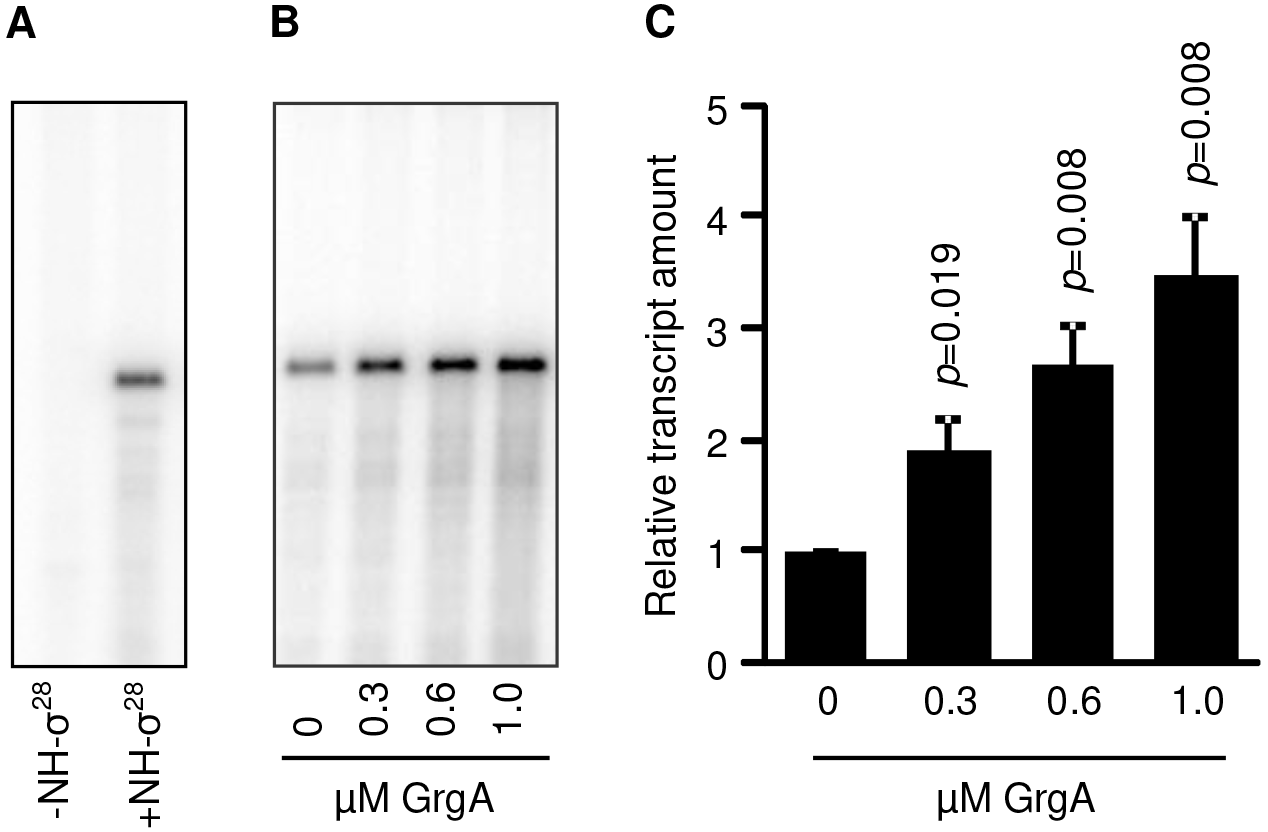
GrgA stimulates σ^28^-dependent transcription using *C. trachomatis* RNAP. (A) Transcription from the *C. trachomatis hct*B promoter in the pMT1212 report plasmid is dependent on the addition of NH-σ^28^ to the chlamydial RNAP. (B) Gel image showing dose-dependent stimulation of transcription from the *hct*B promoterby GrgA. (C) Averages and SDs for three independent measurements are shown.

### Residues 138-165 in GrgA are required for binding both σ^28^ and DNA, and for activating σ^28^-dependent transcription

A series of His-tagged GrgA deletion mutants (Fig. S3A, B) were tested for the effects on transcription from the *hct*B promoter (Fig. 3A). Noticeably, GrgAΔ 114-165 was completely defective in activating transcription from the σ^28^-dependent promoter, whereas GrgAΔ1-64 also demonstrated a significant 50% loss of transcription activation activity (Fig. 3A). Deletion of other regions (65-113, 166-266 and 207-288) from GrgA had either no or minimal effects on the transcription activation (Fig. 3A).

**Fig. 3.**
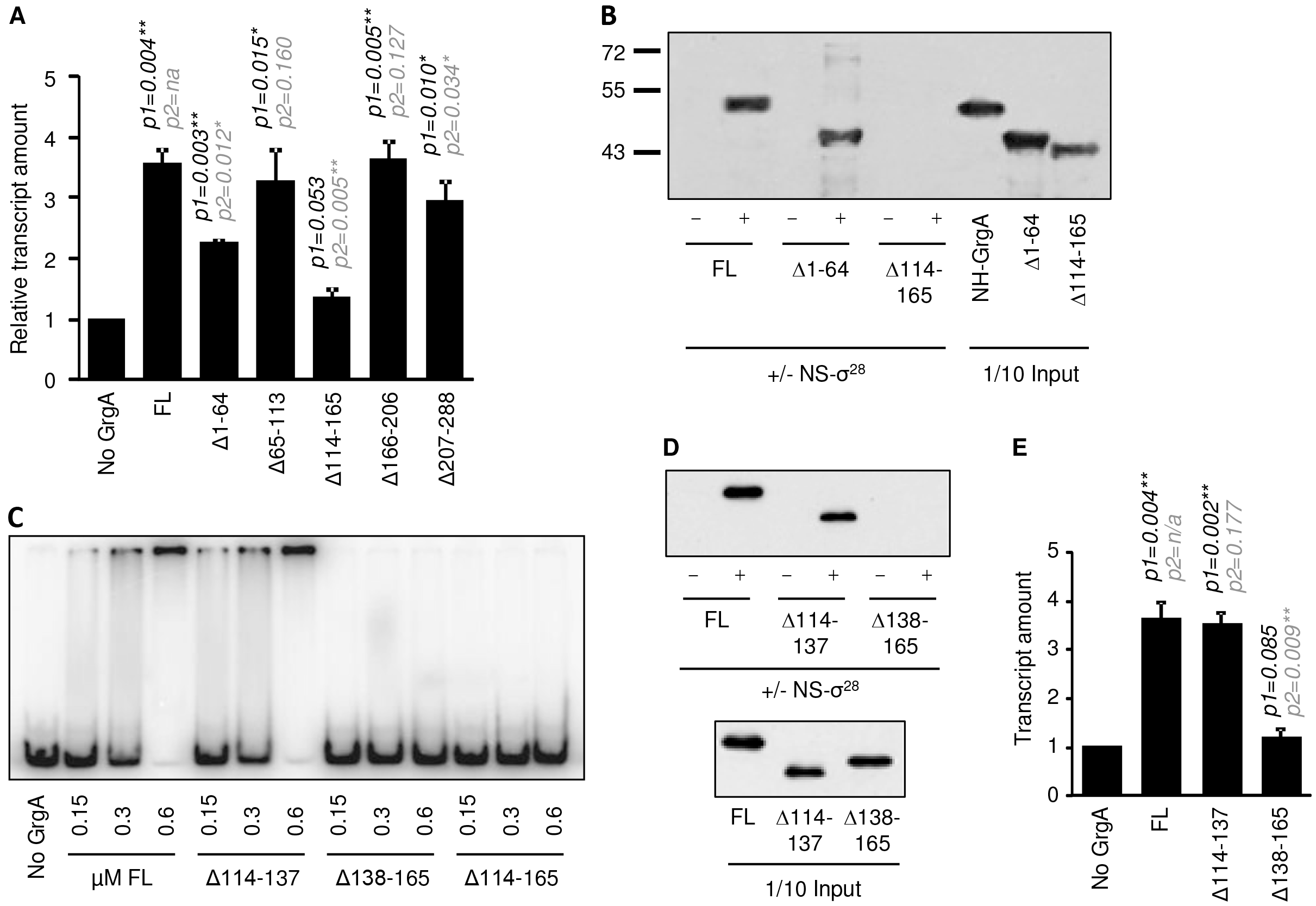
Residues 138-165 in GrgA are required for binding both σ^28^ and DNA, and for activating σ^28^- dependent transcription. (A) Efficient stimulation of transcription from the hctB promoter requires the N-terminal residues 1-64 and the middle region (114-165). 1 full length GrgA (FL) or indicated deletion mutant was used per assay. *p1* is the p value between basal transcription activity (without GrgA) and activity with GrgA (FL or mutant); *p2* is the *p* value between FL and the deletionmutant (paired *t* tests of three independent experiments); na, not applicable. (B) Pull-down of NH-GrgA full length (FL) and Δ1-64 but not Δ 114-165 by StrepTactin bead-immobilized NS-σ^28^. GrgA and deletion mutants were detected by western blotting using a mouse anti-GrgA antibody because the anti-His used in (A & B) does not recognize Δ1-64 (18). (C) Δ138-165 but not Δ114-137 is defective in DNA-binding like Δ114-165. Electrophoresis mobility shift assays (EMSA) were performed using a radiolabeled DNA fragment carrying sequences extending from position –144 to +52 of the *def*A gene (18) in the presence of the indicated concentrations of wild-type GrgA or the indicated GrgA mutant. (D) Δ138-165 but not Δ114-137 is defective in σ^28^-binding. Protein pull-down and detection were performed as described in B, with the exception of detection via anti-His. (E) Δ138-165 but not NH-GrgAΔ114-137 is fully defective in activating transcription from the *hct*B promoter. See Fig. 2D legend for experimental and statistical information.

Our previous studies have shown that deletion of residues 114-165 disables GrgA’s DNA-binding, leading to loss of stimulation of transcription from σ^66^-dependent promoters, whereas removal of residues 1-64 disables σ^66^-binding, also causing defect in activating σ^66^-dependent transcription (18). Therefore, the results in Fig. 3A suggests that 1) DNA-binding is also required for σ^28^-dependent transcription, and 2) the N-terminal σ^66^-interacting region may interact with σ^28^ as well. Surprisingly, pull-down assays demonstrated that GrgAΔ114-165 is completely defective in σ^28^-binding, whereas GrgAΔ1-64 appeared to have only a slightly decreased σ^28^-binding activity (Fig. 3B).

We performed a series of deletions within the 114-165 region to define the elements required for interacting with either DNA or σ^28^. Since residues 114-138 are predicted to have coiled and stranded structures, whereas residues 139-158 are rich in positively charged lysine and aspartate, and are predicted to form a helix (Fig. S4), we expected GrgAΔ114-137 but not GrgAΔ138-165 to retain DNA-binding activity. EMSA confirmed this prediction (Fig. 3C). Interestingly, GrgAΔ114-137 but not GrgAΔ138-165 also retained σ^28^-binding activity as well (Fig. 3D). Not surprisingly, GrgAΔ114-137 but not GrgAΔ138-165 retained the capacity to activate σ^28^-dependent transcription (Fig. 3E). Additional and extensive deletion mutagenesis and functional analyses for the region of residues 138-165 failed to 1) separate residues required for σ^28^-binding from residues required for DNA-binding (Fig. S5 & S6), and 2) define a smaller region fully required for binding either σ^28^ or DNA (Fig. S5 & S6). These studies suggest that σ^28^ and DNA bind to the same region in GrgA, and further confirm that σ^28^- and DNA-binding are required for activation of σ^28^-dependent transcription (Fig. S7).

### Residues 1-64 in GrgA contribute to σ^28^-binding

Transcription assays showed a 50% loss of activity in activating σ^28^-dependent transcription in the GrgAΔ1-64 mutant (Fig. 3A). We used the BLItz system (20) to confirm decreased σ^28^-binding activity in Δ1-64. Representative BLItz recordings of binding experiments using full length NH-GrgA or deletion mutants as ligands and NS-σ^28^ as analyte are shown in Fig. S8A-D, and kinetic parameters are provided in Table 2. Compared to the full length NH-GrgA, NS-σ^28^ revealed a moderately slowed association with and a moderately accelerated dissociation from the NH-GrgAΔ1-64, as indicated by a nearly 3-fold decrease in the *k*_a_ and a 2-fold increase in the *k*_d_ (Table 2). These changes resulted in a highly significant 4.8-fold increase in the *K*_D_ value. On the other hand, NH-GrgAΔ114-137, which retains the activity to activate σ^28^-dependent transcription (Fig. 3E), demonstrated no changes in kinetic parameters for interaction with NS-σ^28^ (Table 2). In contrast to NH-GrgAΔ114-137, NH-GrgAΔ138-165 immediately and completely dissociated from NS-σ^28^ upon wash (Fig. S8D), leading to a 1.5 X 10^6^ times higher *k*_d_ and a 3.1 X 10^6^ times higher *K*_D_ (Table 2), which are fully consistent with pull-down data (Fig. 3D). Furthermore, unlike NS-σ^28^, which retained a low affinity for NH-GrgAΔ1-64 (relative to full length NH-GrgA), CS-σ^66^ quickly and completely dissociated from NH-GrgAΔ1-64 upon wash (Fig. S8E), which is consistent with pull-down data previously reported (18). Taken together, the BLItz data in Fig. S8 and Table 2 indicate that the decreased affinity with NS-σ^28^ in NH-GrgAΔ1-64 is responsible for the partial loss of activity in activating σ^28^-dependent transcription (Fig 3A).

**Table 2.**
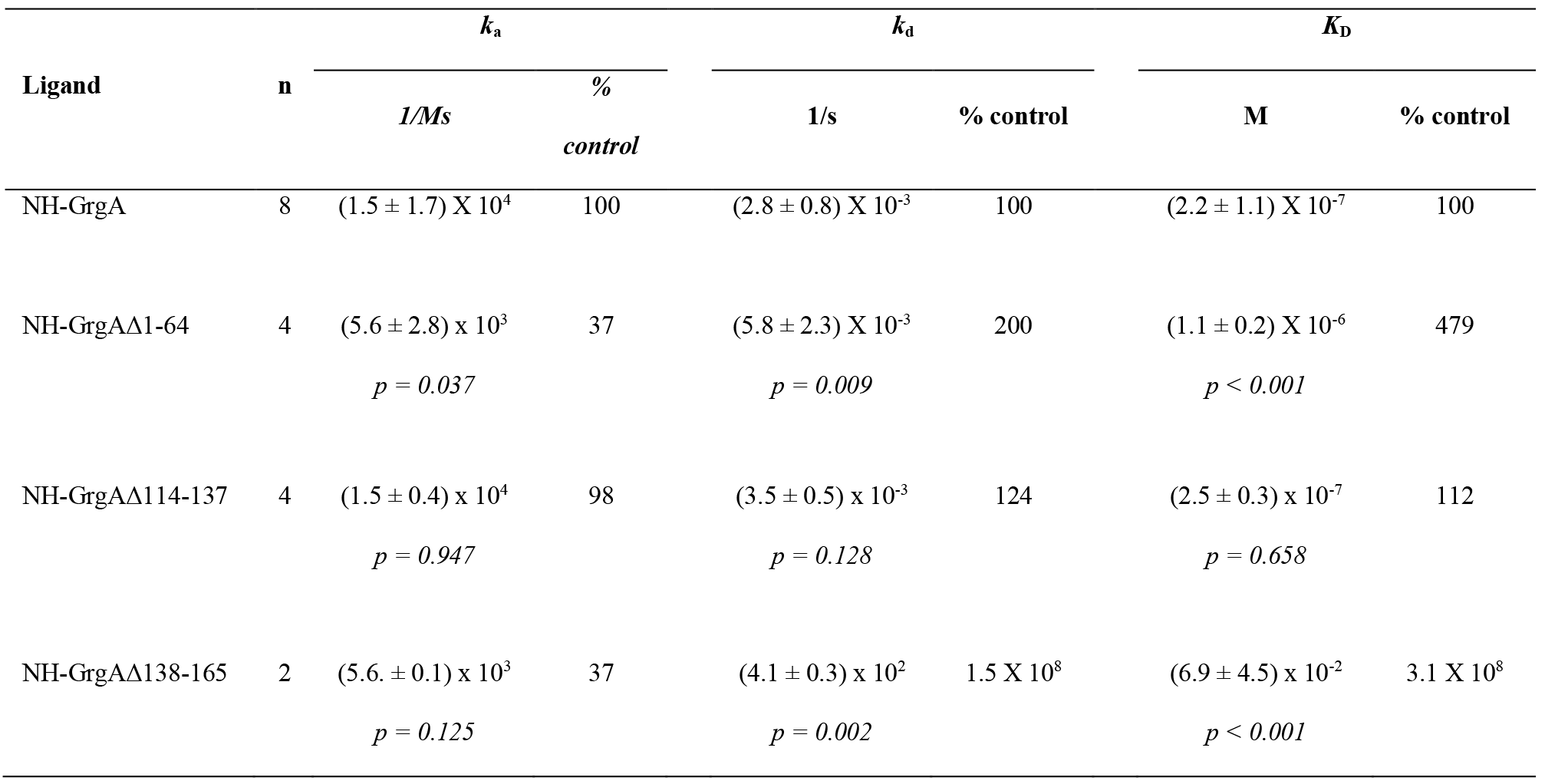
Deletion of amino acids 1-64 from GrgA negatively affects σ^28^-binding. Indicated His-tagged proteins were immobilized on His biosensors as ligands for the NS-σ^28^ analyte in BLItz assays. Representative graphs of BLItz recordings are shown in Fig. S8. See Table 1 for information regarding kinetic parameters and statistics.

### The N-terminus of σ^28^ is most critical for binding GrgA

We constructed NH-σ^28^ variants with deletion of the N-terminal leader sequence, σ factor region 2, 3 or 4 (unlike the housekeeping σ factor, σ^28^ does not contain region 1) (Fig. 4A). All deletion mutants (i.e., NH-σ^28^ΔNL, NH-σ^28^ΔR2, NH-σ^28^ΔR3 and NH-σ^28^ΔR4) were expressed in *E. coli* (Fig. 4B). Noticeably, in pull-down assays, NS-GrgA completely failed to pull down the NH-σ^28^ΔNL and NH-σ^28^ΔR2 mutants, and pulled down only small amounts of NH-σ^28^ΔR3 and NH-σ^28^ΔR4, compared to full length NH-σ^28^ (Fig. 4C).

**Fig. 4.**
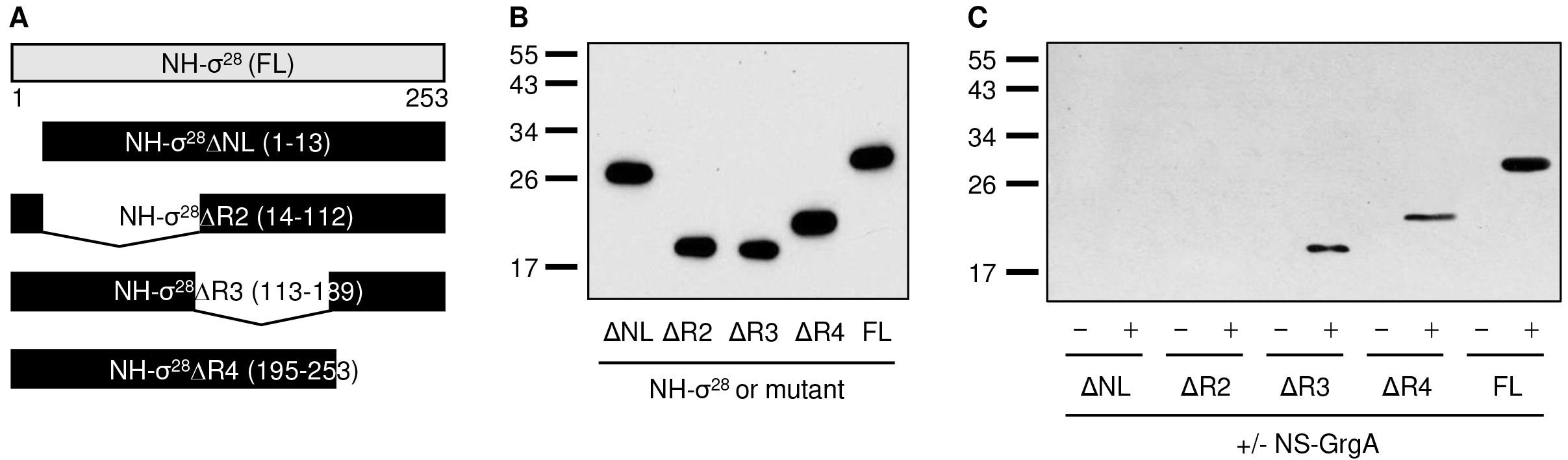
The N-terminal lead sequence and region 2 of σ^28^ are requiredfor interaction with GrgA. (A) Schematic of σ^28^ and mutants lacking the indicated regions. (B) Western blot showing expression of purified NH-σ^28^ and mutants. (C) Precipitation of NH-σ^28^ and mutants by StrepTactin-immobilized NS-GrgA. Shown is a western blot detecting NH-σ^28^ or mutant. All proteins were resolved via SDS-PAGE and detected using an anti-His antibody.

In BLItz assays, the rate of association with NH-GrgA varied greatly among the σ^28^ deletion mutants. Whereas NH-σ^28^ΔNL had essentially the same *k*a as full length NH-σ^28^, NH-σ^28^ΔR2 displayed a significant 2-fold reduction in *k*_a_. In contrast, NH-σ^28^ΔR3 and NH-σ^28^ΔR4 showed a 3.5-fold increase and a 50% increase in *k*_a_, respectively (Table 3). All mutants demonstrated dramatic increases (10-383 fold) in the *k*_d_ (Table 3). Consequently, NH-σ^28^ΔNL and NH-σ^28^ΔR2 had 345- and 177-fold higher *k*_D_ value, respectively, whereas NH-σ^28^ΔR3 and NH-σ^28^ΔR4 both demonstrated 7 fold higher *K*_D_ values (Table 3). Representative graphs of BLItz recordings are shown in Fig. S9. Taken together, both the pull-down (Fig. 4) and BLI data (Table 3) indicate that the N-terminus of σ^28^ (i.e., NL and R2) interacts with GrgA while R3 and R4 stabilize the GrgA-σ^28^ binding.

**Table 3.**
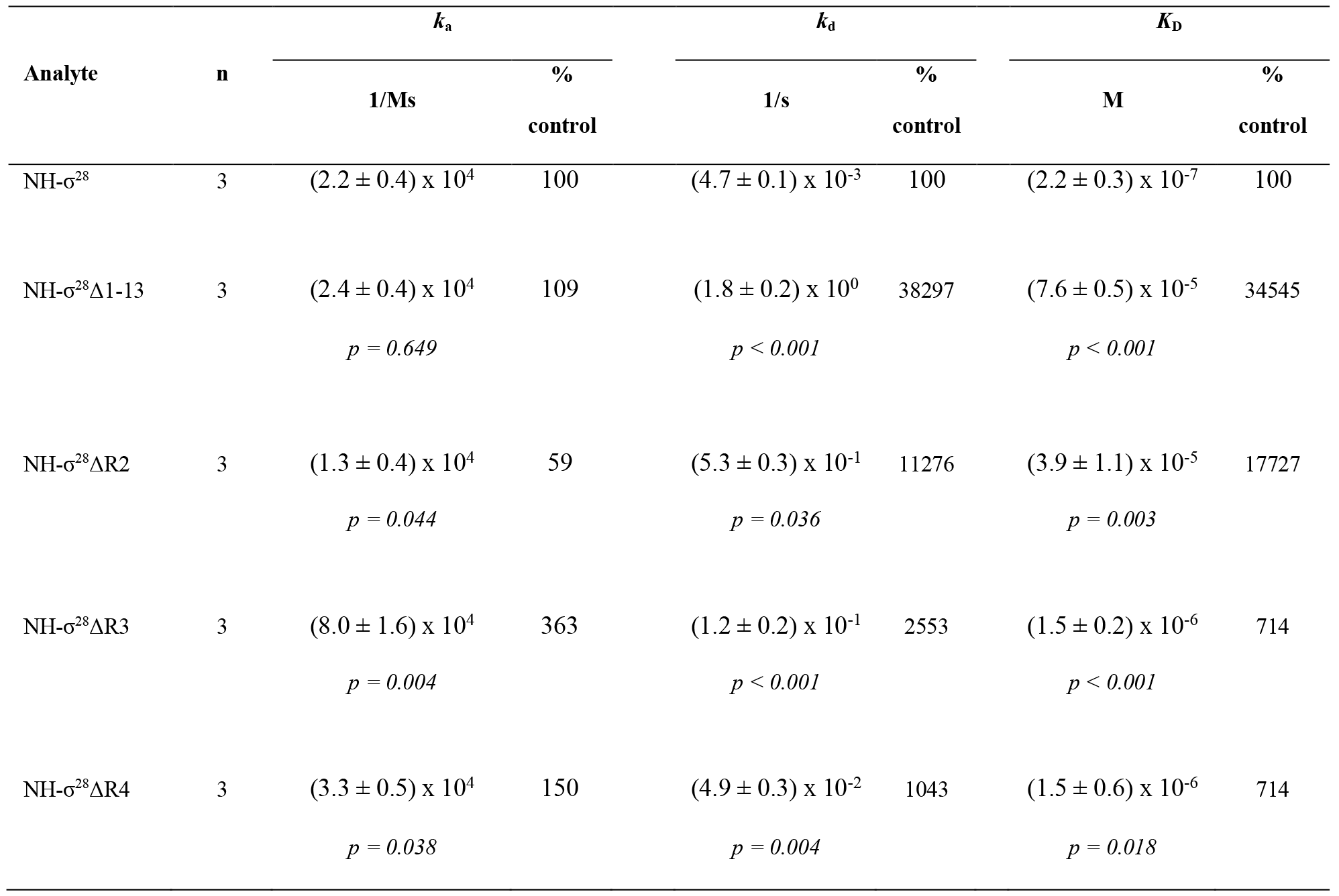
The N-terminal leading sequence and region 2 of σ^28^ are required forGrgA binding. Biotinylated NH-GrgA was immobilized on streptavidin biosensors as the ligand for indicated analytes in BLItz assays. Representative graphs of recordings are shown in Fig. S9. See Table 1 for information regarding kinetic parameters and statistics.

## DISCUSSION

Although GrgA was first identified as a transcription activator for σ^66^-dependent genes (18), the present study has demonstrated that GrgA potentially stimulates expression of σ^28^-dependent genes. Transcription of chlamydial genes is temporally controlled during the developmental cycle (11, 12, 17, 22, 23). Whereas σ^66^ is involved in transcription of most *C. trachomatis* genes, some late promoters are recognized by σ^28^ (17). Microarray studies have shown that synthesis of the σ^28^ mRNA temporally falls behind the σ^66^ mRNA (11, 12). Thus, it would be safe to assume that GrgA primarily activates σ^66^-dependent genes in earlier developmental stages.

Whether or not GrgA also regulates expression of σ^28^-dependent genes during later developmental stages likely depends on the expression levels of GrgA,σ^28^ and σ^66^. If GrgA is limited, σ^28^ would have to be present at significantly higher concentrations than σ^66^ to effectively compete for GrgA. However, quantitative whole proteomic mass spectrometry analyses detected higher levels of GrgA relative to σ^66^ in both EBs and RBs purified from the midcycle (24) whereas σ^28^ was undetected in either cellular form (24, 25). Thus, GrgA could potentially stimulate transcription from σ^28^-dependent promoters in addition to σ^66^- dependent promoters regardless the molar ratio of the two σ factors. Accurate quantification of GrgA and a factors in different stages of the developmental cycle will help elucidate the role of GrgA in the expression of σ^66^- and/or σ^28^-dependent genes in different developmental stages.

We used both pull-down assays and BLI to analyze the interaction of GrgA with σ^28^ and σ^66^. Clearly, owing to its quantitative nature, BLI offers higher sensitivities than protein pull-down assays in studying protein-protein interaction. This led to the confirmation that decreased affinity for σ^28^ in GrgAΔ1-64, which was ambiguous in pull-down assays, is the most probable cause for a 50% loss of activity in activating σ^28^-dependent transcription.

We have defined a middle region in GrgA (residues 138-165) as a σ^28^- and DNA-binding domain (Fig. 3). Extensive deletion mutagenesis in this region failed to divide it into subdomains that bind either σ^28^ or DNA but not both (Figs. S5 & S6). We speculate that multiple positively-charged residues (K138, K139, R142, R143, K144, K147, K150, K152, K154-156, R159-161 and/or K164) interact with negatively charged DNA whereas multiple negatively-charged residues E141, E145, E149, D153 and/or E165) interact with σ^28^.

GrgA has demonstrated similar but not identical properties in activating σ^66^- and σ^28^-dependent transcription. Apparently, sequence-nonspecific DNA-binding is required for activating both σ^66^- dependent transcription (18) and σ^28^-dependent transcription (Fig. 3A, E & Fig. S7). However, the N-terminal region (residues 1-64) of GrgA has a stronger role in σ^66^-dependent transcription (18) than in σ^28^- dependent transcription (Fig. 3A) because this region is absolutely required for GrgA to interact with σ^66^ (18), but plays only a supportive role in binding σ^28^, which was clearly evident only with BLI (Table 2) but appeared uncertain with pull-down assays (Fig. 3B).

Whereas the major GrgA structural determinants for binding σ^28^ and σ^66^ differ, there is similarity between the GrgA-binding regions in the two σ factors. The GrgA-binding sequence in σ^66^ is the last portion of the non-conserved region immediately upstream of the conserved region 2, whereas the GrgA-binding sequence in σ^28^ also involves the N-terminal non-conserved leader sequence (and the immediately downstream region 2). To the best of our knowledge, GrgA is the only transcription factor that targets non-conserved regions of σ factors (16).

In summary, we have demonstrated that the *Chlamydia*-specific GrgA can activate both σ^66^-dependent transcription and σ^28^-dependent transcription *in vitro.* Current knowledge suggests that GrgA primarily activates σ^66^-dependent genes during earlier developmental stages. However, whether or not GrgA also regulates expression of σ^28^-dependent genes during later developmental stages likely depends on the expression levels of GrgA, σ^28^ and σ^66^ because GrgA has a lower affinity for σ^28^ than σ^66^. To date, GrgA remains the only transcription factor that targets non-conserved regions of σ factors (16).

## MATERIALS AND METHODS

### Reagents

All DNA primers were custom-synthesized at Sigma Aldrich. The QuikChange Site-Directed Mutagenesis Kit, BL21(DE3) ArcticExpress *E. coli* competent cells were purchased from Agilent Technologies. Q5 Site-Directed Mutagenesis Kit, and deoxynucleotides were purchased from New England BioLabs. Isopropyl β-D-1-thiogalactopyranoside (IPTG) was purchased from Gold Biotechnologies. TALON Metal Affinity Resin was purchased from Takara. The StrepTactin superflow high capacity resin and D-desthiobiotin were purchased from IBA Life Sciences. Coomassie Brilliant Blue G-250 Dye, mouse monoclonal anti-Histidine antibody (H1029), goat anti-mouse horseradish-perodixase-conjugated antibody (A4416), and EZ-Link Sulfo-NHS-LC-Biotin were purchased from Sigma Aldrich. SuperSignal West Pico PLUS Chemiluminescent Substrate was purchased from ThermoFisher Scientific. Dip and Read Ni-NTA (NTA) biosensors were purchased from Pall ForteBio. The HNE Buffer contained 50 mM HEPES (pH 7.4), 300 mM NaCl, and 1 mM EDTA. The HNEG Buffer contained 50 mM HEPES (pH 7.4), 300 mM NaCl, 1 mM EDTA, and 6M Guanidine HCl. The TNE Buffer contained 25 mM Tris (pH 8.0), 150 mM NaCl, and 1 mM EDTA. The Protein Storage (PS) Buffer contained 25 mM Tris-HCl (pH 8.0), 150 mM NaCl, 0.1 mM EDTA, 10 mM MgCl_2_, 0.1 mM DTT, and 30% glycerol (w/v). The BLItz Buffer contained 25 mMTris-HCl (pH 8.0), 150 mM NaCl, 0.1 mM EDTA, 10 mM MgCl_2_, and 0.1 mM DTT.

### Vectors

Plasmids for expressing His- or Strep-tagged GrgA, σ^66^, σ^28^, and their mutants are listed in the Table S1. Sequences of primers used for constructing expression plasmids (GrgA deletion mutants, NS-σ^28^)and a DNA fragment for EMSA assays are available upon request. Sequence authenticities of cloned genes and epitope tags in the final vectors were confirmed using Sanger’s DNA sequencing service provided by GenScript Biotech Corporation.

### Expression of recombinant proteins and preparation of cell extract for purification

BL21(DE3) ArcticExpress *E. coli* cells transformed with a plasmid for expressing an epitope-tagged chlamydial protein (GrgA, σ^28^, σ^66^ or their mutant) (Table S1) were cultured in the presence of 1 mM IPTG overnight at 15 °C in a shaker. Cells were collected by centrifugation and resuspended in one of the following buffers: HNE buffer (for purification of native His-tagged proteins), HNEG buffer (for purification of denatured His-tagged proteins), or TNE buffer (for purification of native Strep-tagged proteins). The cells were disrupted using a French Press. The cell extract was subjected to high-speed (20,000g) centrifugation at 4 °C for 30 minutes. Supernatant was collected and used for protein purification.

### Purification of Strep-tagged proteins

Strep-tagged GrgA and σ factors were purified as previously described (19). The supernatant of centrifuged cell lysate was incubated with the StrepTactin superflow high capacity resin on a Nutator for 1 h at 4 °C. The resin was packed onto a column and washed with 30 column volumes of the TNE Buffer, and then eluted with the TNE Buffer containing 2.5 mM D-desthiobiotin. The elution was collected in 10 fractions. Protein in the fractions was examined following SDS-PAGE and Coomassie-Blue staining. Fractions with high purity and concentration were pooled and dialyzed overnight against the PS Buffer at 4 °C, and then stored in aliquots at −80 °C.

### Purification of His-tagged proteins

The supernatant of centrifuged cell lysate was incubated with the TALON metal affinity resin on a Nutator for 1 hour at 4 °C. The resin incubated with non-denatured cell extract was packed onto a column, washed with 30 column volumes of HNE Buffer containing 1% NP-40, and eluted with the HNE Buffer containing 250 mM imidazole. The resin incubated with denatured cell extract was packed onto a column, washed with 30 column volumes of HNEG Buffer, and eluted with HNEG Buffer containing 250 mM imidazole. Examination of protein purity, dialysis and storage were carried out in the same manner as for purified Strep-tagged proteins (18).

### In vitro transcription assay

*In vitro* transcription of σ^28^-dependent promoter was performed as previously described (18). The assay in a total volume of 30 μl contained 200 ng supercoiled plasmid DNA, 50 mM potassium acetate, 8.1mM magnesium acetate, 50 mM Tris acetate (pH8.0), 27 mM ammonium acetate, 1 mM DTT, 3.5% (wt/vol) poly-ethylene glycol (average molecular weight, 8,000), 330 μM ATP, 330 μM UTP,1 μM CTP, 0. 2 μM [α-^32^P]CTP (3,000 Ci/mmol), 100 μM 3’-O-methyl-GTP, 20 units of RNasin, RNAP, and indicated amount of GrgA or GrgA mutant. The reactions using cRNAP and σ^28^ contained 1.0 μL purified cRNAP and 30nM His-tagged σ^28^, purified by procedures involving denaturing and refolding as described above. For reactions using eCore and σ^28^, their concentrations were 5 nM and 30 nM, respectively. The reaction was allowed to pursue at 37 °C for 40 min and terminated by the addition of 70 μL of 2.86 M ammonium acetate containing 4mg of glycogen. After ethanol precipitation, ^32^P-labeled RNA was resolved by 8M urea-6% polyacrylamide gel electrophoresis, and quantified with a Storm Phosphorimager and the ImageQuant software. Relative amounts of transcripts were presented with that of the control reaction set as 1 unit. Data shown in bar graphs represent averages ± SDs from three or more independent experiments. Pairwise, two-tailed Student *t* tests were used to compare data.

### Electrophoresis mobility shift assay (EMSA)

GrgA-DNA interaction was determined by EMSA as described previously (18). ^32^P-labeled DNA fragment containing the *C. trachomatis def*A promoter (26) was amplified using a ^32^P-labeled 5’ primer and an unlabeled 3’ primer (Table S2) and purified with a Qiagen column. The GrgA-DNA binding reaction was performed in a total volume of 10 μL, containing 10 nM promoter fragment, an indicated amount of NH-GrgA, 1 mM potassium acetate, 8.1 mM magnesium acetate, 50 mM Tris acetate (pH 8.0), 27 mM ammonium acetate, 1 mM DTT, and 3.5% (wt/vol) polyethylene glycol (average molecular weight, 8,000). After mixing for 1 h at 4 °C, the binding mixture was resolved by 6% non-denaturing polyacrylamide gel. Free and GrgA-bound DNA fragments were visualized on a Storm Phosphorimager (Molecular Dynamics).

### Pull-down assays

20 μL of StrepTactin superflow high-capacity resin was washed twice with the HNE Buffer and incubated with 50 μL of Strep-tagged cell extract or purified protein on a Nutator at 4°C for 1 h. The resin was washed three times with HNE Buffer containing 1% NP-40, and then incubated with 5 μg of a purified His-tagged protein (or mutant) on a Nutator at 4 °C for 1 h. After 3 washes with the HNE Buffer containing 1% NP-40 and a final wash with PBS, the resin was eluted using SDS-PAGE sample buffer. All protein was resolved via SDS-PAGE and detected by either Coomassie blue staining or western blotting using a monoclonal mouse anti-His or a polyclonal mouse anti-GrgA primary antibody and HRP-conjugated goat anti-mouse secondary antibody.

### Preparation of biotinylated protein

Purified NH-GrgA was dialyzed against PBS to remove Tris and then incubated with 10 mM EZ-Link Sulfo-NHS-LC-Biotin for 2 hours at 4 °C. Excess biotin was removed via two-step dialysis, initially against PBS and subsequently against the PS buffer.

### Bio-layer interferometry assay

An NTA His or streptavidin biosensor was subjected to initial hydration in BLItz Buffer for 10 minutes before being loaded onto the ForteBio BLItz machine and washed with BLItz Buffer for 30 seconds to obtain a baseline reading. The His biosensor was then incubated with 4 μL of a His-tagged ligand for 240 seconds. The concentration of ligand ranged from 1-20 μM, which all saturated the His-binding sites on the biosensor. Alternatively, the streptavidin biosensor was incubated with 4 μL of a biotinylated ligand (NH-GrgA, 10 μM, which was sufficient to saturate the binding sites on the biosensor) for 240 seconds. After a brief wash with BLItz Buffer for 30 seconds to remove excess protein, the biosensor was incubated with 4 μL of an analyte (purified Strep-tagged protein for the His biosensor or NH-σ^28^ for the streptavidin biosensor) for 120 seconds to measure association of the ligand-analyte complex. Subsequently, the biosensor was washed with BLItz Buffer for 120 seconds to measure disassociation of the ligand-analyte complex. All BLItz recordings were subsequently fit to a 1:1 binding model using the BLItz Pro software (version 1.1.0.31), which generated the association rate constant (*k*_a_), disassociation rate constant (*k*_d_), and disassociation equilibrium constant (*K*_D_) for each interaction.

## ACKNOWLEDGEMENTS

We thank Ming Tan (University of California Irvine) for pMT1212. This research was supported by National Institutes of Health (Grant # AI122034 to HF, GM118059 to BEN), New Jersey Health Foundation (Grant # PC 20-18 to HF) and Natural Sciences Foundation of China (Grant # 31370209 and 31400165 to XB).

